# Tobacco Hornworm (*Manduca sexta*) caterpillars as a novel host model for the study of fungal virulence and drug efficacy

**DOI:** 10.1101/693226

**Authors:** Naomi Lyons, Isabel Softley, Andrew Balfour, Carolyn Williamson, Heath E. O’Brien, Amol C. Shetty, Vincent M. Bruno, Stephanie Diezmann

## Abstract

The two leading yeast pathogens of humans, *Candida albicans* and *Cryptococcus neoformans*, cause systemic infections in >1.4 million patients world-wide with mortality rates approaching 75%. It is thus imperative to study fungal virulence mechanisms, efficacy of antifungal drugs, and host response pathways. While this is commonly done in mammalian models, which are afflicted by ethical and practical concerns, invertebrate models, such as wax moth larvae and nematodes have been introduced over the last two decades. To complement existing invertebrate host models, we developed fifth instar caterpillars of the Tobacco Hornworm moth *Manduca sexta* as a novel host model. These caterpillars can be maintained at 37°C, are suitable for injections with defined amounts of yeast cells, and are susceptible to the most threatening yeast pathogens, including *C. albicans*, *C. neoformans*, *C. auris*, and *C. glabrata*. Importantly, fungal burden can be assessed daily throughout the course of infection in a single caterpillar’s faeces and haemolymph. Infected caterpillars can be rescued by treatment with antifungal drugs. Notably, these animals are large enough for weight to provide a reliable and reproducible measure of fungal disease. *M. sexta* caterpillars combine a suite of parameters that make them suitable for the study of fungal virulence.

## Introduction

Fungal infections pose a serious threat to human health and well-being world-wide. Each year, as many, if not more patients, die of fungal infections than of malaria or tuberculosis^1^. The leading yeast pathogens, *Candida albicans* and *Cryptococcus neoformans*, account for ~1,400,000 life-threatening infections world-wide with mortality rates of up to 70%^1,2^. Candidaemia, most frequently caused by *C. albicans*, is the fourth most common cause of nosocomial blood stream infections, only surpassed by infections with Staphylococci and *Enterococcus* spp. Disturbingly, candidaemia incidence rates are on the rise. Within less than ten years, they increased by 36%^3^. Although cryptococcosis incidence rates are on the decline in North America, cryptococcosis as an AIDS-defining illness is responsible for 15% of all AIDS-related deaths world-wide^3,4^. This dire situation is further confounded by the emergence of drug resistant yeast species, such as *C. glabrata* and *C. auris*. Patients at risk of developing invasive candidaemia are often prophylactically treated with fluconazole and echinocandins as a first line defence strategy^5^. Yet, *C. glabrata*, the most common non-*albicans Candida* species associated with nosocomial blood stream infections^6^, is intrinsically less susceptible to azole drugs and acquires resistance to echinocandins rapidly^7^. The rapid global spread of multi-drug resistant *C. auris* has further exacerbated the threat posed by fungal infections. *C. auris* was first reported in 2009 in Japan^8^. In 2015, *C. auris* arrived in Europe causing an outbreak involving 72 patients in a cardio-thoracic hospital in London^9^. *C. auris* outbreaks have been reported from South Korea, India, Spain, Columbia, Switzerland, Germany, Israel, Kuwait, and Oman^10^. Most concerningly, up to 25% of *C. auris* isolates are multi-drug resistant, with some strains being resistant to three of the four drug classes available for the treatment of systemic candidaemia. In addition to the unacceptably high burden on human health, fungal infections substantially increase health care costs. Treatment of candidaemia requires extended hospitalisation, resulting in additional costs of up to $45,000 in adult patients or up to $119,000 in paediatric patients^11^.

It is thus imperative to investigate fungal virulence and host response mechanisms. This is traditionally done in mammalian models. The most frequently deployed models include the mouse tail vein infection model for systemic candidaemia, the mouse gastrointestinal infection model of candidaemia, the mouse *Candida* vaginitis model^12^, the mouse inhalation model of cryptococcosis^13^, the rabbit chronic cryptococcal meningitis model^14^, and the rabbit *Candida* keratitis model^15^. While mammalian models combine a number of features that make them particularly amenable for the study of fungal diseases, such as susceptibility, availability of knock-out mutants, and comparable histology to human disease, using mammals is ethically controversial, economically challenging, and requires extensive board certifications and documentations.

In an effort to reduce the usage of mammalian hosts, alternative invertebrate models have been developed and successfully used over the past two decades. The most commonly employed invertebrate species include the nematode *Caenorhabditis elegans*, the fly *Drosophila melanogaster*, and larvae of the Greater Wax moth *Galleria mellonella*. All three can be easily maintained in the laboratory, are much less expensive than mice or rabbits, and have been used for the study of diverse yeast pathogens, such as *C. neoformans*^16,17^, *C. albicans*^17–19^, *C. parapsilosis*^19–21^, *C. glabrata*^20,22^. Of note, invertebrate models differ in their applicability and the best suitable model should be carefully selected^23^. Unlike mammalian models, these invertebrates do not have adaptive immunity but all share components of the innate immune system^24,25^, some of which are conserved with mammals. This includes the Toll-like receptors found in the fly^26^ and the homolog of the MKK3/6 kinase in the nematode^27^. Ironically, it is the Toll-like receptors that protect flies from infections with *C. neoformans*^28^, *C. albicans*^29^, and *C. glabrata*^30^ and the MKK3/6 homolog SEK-1 protects the nematode from bacterial invaders^27^. Thus, to increase susceptibility of flies and nematodes to fungal pathogens, Toll and *sek-1*^31^ mutants are required. A key limitation for the study of human pathogens, is the inability of nematodes and flies to survive human body temperature. Only *Galleria* can withstand 37°C^17^. Due to their long-standing history as eukaryotic models, well-curated genomes and genome databases exist for the nematode and the fly. Although the *Galleria* genome has been announced very recently^32^, detailed analyses and annotations are still outstanding. Delivery of an exact inoculum of fungal cells is crucial when comparing mutant and wild type yeast strains, neither fly nor nematode permit routine direct injection. *Galleria* can be directly injected with a defined cell number. Extensive stock collections provide fly and nematode strains, while until recently, *Galleria* larvae had to be purchased from fishing shops. Now, UK-based TruLarv is selling research grade larvae.

Tobacco Hornworms are most commonly encountered in the southern United States, where they feed on solanaceous plants and are thus considered a plant pest. As an insect model with a long history in research, *Manduca sexta* has yielded important insights into flight mechanisms, nicotine resistance, hormonal regulation of development, metamorphosis, antimicrobial defences, and bacterial pathogenesis. *M. sexta* laboratory stocks have been derived from animals collected in North Carolina, USA^33^ and have been maintained in laboratories on both sides of the Atlantic for several decades. *M. sexta*’s research portfolio includes a draft genome sequence that has been complemented with tissue-specific transcriptomic analyses^34^, numerous successful applications of RNAi^35–39^, and protocols for the efficient extraction of hemocytes for down-stream analyses^40^ of the animal’s innate immunity^41^. Despite its versatility, *M. sexta* has yet to be explored for its suitability as a host model for fungal infections.

Here, we aimed to establish *M. sexta* as a novel model host for the study of fungal virulence. Inbred animals from the University of Bath’s research colony were tested for their ability to live at 37°C, their susceptibility to different yeast species, and the reproducibility of *C. albicans* mutant phenotypes obtained in mice virulence studies. Indeed, the caterpillars grow at 37°C while maintaining susceptibility to the leading yeast pathogens *C. albicans*, *C. neoformans*, and emergent *C. auris*. Specific *C. albicans* mutants are just as attenuated in their virulence in *M. sexta* as they are in mice. To expand *M. sexta’s* applicability as a host model, we developed an infection protocol that permits screening of fungal burden throughout the course of infection in a single animal and uses weight as a proxy measure for virulence in addition to survival. *M. sexta* can furthermore be used to test efficacy of common antifungal drugs and to query the host transcriptional response to systemic yeast infections. Our results define *M. sexta* characteristics that recommend these caterpillars as a non-mammalian host model for the study of fungal virulence with susceptibility to different yeast species.

## Materials and Methods

### Origin of the Bath colony of *Manduca sexta*

The University of Bath colony has been in continuous culture since 1978 without the addition of animals from elsewhere. Bath’s genetic stock was derived from animals from the Truman-Riddiford laboratories at the University of Washington in Seattle, USA. Their animals date back to the ones originally collected in North Carolina in 1976^33^.

### Caterpillar maintenance and yeast culture conditions

*M. sexta* caterpillars were reared to fifth instar under standardised conditions. They were maintained in 125 ml disposable cups (Sarstedt Ltd., Cat. No. 75.1335) on a wheat germ-based diet (Appendix ‘Food preparation’) at a constant temperature of 25°C with 50% humidity and 12 hours light/dark cycles. Three days prior to fungal inoculations, animals were shifted to a formaldehyde-free diet as the compound is toxic to non-methylotrophic yeast.

For infection assays, yeasts were grown overnight in 50 ml YPD (1% yeast extract, 2% peptone, 2% dextrose) and cells harvested by centrifugation for 3 minutes at 3,000 rpm. The cell pellet was washed twice with 1x phosphate buffered saline (PBS) and suspended in 5 ml 1x PBS. Cells were counted and numbers adjusted as indicated. *C. albicans* YSD85 (Table 1) cells were heat-inactivated by incubation at 65°C for 20 minutes. For long-term storage, yeast isolates were cryo-archived at −80°C in 25% glycerol.

**Table 1:** Yeast strains used in this study.

| Strain ID | Strain Name/genotype | Species | Source |
| --- | --- | --- | --- |
| YSD89 | SN95 | *C. albicans* |  |
| YSD85 | SC5314 | *C. albicans* |  |
| YSD883 | *hog1Δ/Δ* | *C. albicans* | This study |
| YSD87 | BWP17 | *C. albicans* | 76 |
| YSD190 | VIC84 (*cka2Δ/Δ*) | *C. albicans* | 77 |
| YSD192 | VIC93 (*cka2/cka2::CKA2*) | *C. albicans* | 77 |
| YSD302 | CAS12 (*ahr1Δ/Δ*) | *C. albicans* | 52 |
| YSD303 | CAS13 (*ahr1/ahr1::AHR1*) | *C. albicans* | 52 |
| YSD304 | RM1000 | *C. albicans* | 78 |
| YSD305 | JC52 (*hog1/hog1::HOG1*) | *C. albicans* | 78 |
| YSD306 | JC50 (*hog1Δ/Δ*) | *C. albicans* | 78 |
| YSD790 | YPS143 | *S. cerevisiae* | 79 |
| YSD1448 | NCYC2580 | *M. pulcherrima* | 80 |
| YSD465 | 2001 | *C. glabrata* | 81 |
| YSD1454 | TA004-14 | *C. auris* | 82 |
| YSD1028 | H99 | *C. neoformans* | 83 |

### Yeast infections and measurements of fungal burden and drug efficacy

100 μl of washed and number-adjusted yeast suspension were injected into each caterpillar’s distal right proleg with a 30G1/2” needle (BD Microlance) and a 1 ml NORM-JECT syringe. Following injection, each animal’s weight was recorded. Animals were scored for survival and weight once daily for three or four days post infection. During the course of the experiment, animals were kept on a 12-hour light/dark cycle at the temperature indicated and on their regular diet.

To measure fungal burden in caterpillar faeces and haemolymph, six animals were injected with 100 μl of either 1x PBS or 10^6^ cells of the wild type YSD89 or the *hog1* mutant strain YSD883 and kept at 37°C. On day 1, two animals were selected from each group. These animals were weighted and their haemolymph and faeces collected daily throughout the course of infection. To collect haemolymph, animals were first kept on ice for 15 minutes. The ‘horn’ was then surface sterilised with 70% ethanol and its top 1-2 mm clipped with a pair of micro scissors. Haemolymph was collected in a pre-chilled 1.5 ml Eppendorf tube and cooled immediately to reduce polymerisation and melanisation. One faecal pellet was collected daily with sterile forceps, weighted and suspended in 500 μl 1x PBS. Prior to diluting, the suspension was thoroughly vortexed for 10 seconds, and centrifuged for 5 seconds using a table top centrifuge to separate faecal matter. To quantify fungal burden, haemolymph and faecal samples were plated either directly onto YPD-agar with Kanamycin 50 μg/ml or in ten-fold serial dilutions. Agar plates were incubated at 30°C for 48 hours and colonies counted.

To assess the efficacy of commonly used antifungal drugs, animals were infected with 10^7^ cells of YSD85 or 100 μl PBS and treated with increasing doses of fluconazole and caspofungin (Sigma Aldrich, Inc.) as indicated. Drugs were injected 30 minutes post-infection with an ethanol-sterilized Hamilton syringe in a total volume of 10 μl per animal. Caterpillars were weighed and scored for survival on the day of injection and the following three days.

### Statistical analyses

Survival plots were made using the survminer R package (https://CRAN.R-project.org/package=survminer), and differences were evaluated using the Kaplan-Meier method. Weight and fungal burden were plotted using ggplot2^42^ and weight differences were evaluated using linear models with day post-inoculation and the interaction between treatment and days post infection as fixed effects and individual as a random effect using nlme (https://CRAN.R-project.org/package=nlme). Upset plots visualising gene expression profiles were made using the UpSetR package^43^. All analyses were conducted in RStudio version 1.1.442.

### RNA sequencing and gene expression analysis

Five animals were injected with 100 μl 1x PBS or 10^6^ yeast cells of the wild type strain SN95 (YSD89) and maintained on a 12-hour light/dark cycle at 37°C for 24 hours. Three animals were then randomly selected for transcriptomic analyses of their midguts. In preparation for midgut extraction, animals were surface sterilized and tissue extracted and washed as described before^34^. Midgut RNA was extracted using TRIzol reagent (Invitrogen, Inc.), quantified using the Agilent 2100 Bioanalyzer system and used to generate RNA-seq libraries (strand-specific, paired end) using the TruSeq RNA sample prep kit (Illumina).

150 nucleotides of the sequence were determined from both ends of each cDNA fragment using the HiSeq 2500 platform (Illumina). An average of 83.79 million read pairs were obtained for each sample (range: 51.4 M to 109.8 M read pairs). Sequencing reads were aligned to the reference *Manduca sexta* whole genome assembly v1.0 (https://i5k.nal.usda.gov/Manduca_sexta) using HISAT2^44^ and alignment files were used to generate read counts for each gene. Statistical analysis of differential gene expression was performed using the DEseq2 package from Bioconductor^45^. A gene was considered differentially expressed if the FDR-value for differential expression was less than 0.10. In order to compare the transcriptional response to *C. albicans* infection by *M. sexta* to that of a mouse, we generated a blast reference database using the peptide sequences for 67,960 proteins from the mouse genome (GRCm38) available from Ensembl (https://useast.ensembl.org). We downloaded the peptide sequences for 27,403 proteins from the *M. sexta* database (OGS v2.0) available from the National Agricultural Library (https://i5k.nal.usda.gov/Manduca_sexta)^34^ provided by the United States Department of Agriculture. We computed peptide sequence similarity between the *M. sexta* proteins and the mouse proteins using the blastp local alignment search tool (ncbi-blast+ v2.8.1)^46^. For each *M. sexta* protein, we had multiple hits that were ranked based on the e-value computed by the blastp search tool. For the list of differentially expressed *M. sexta* genes from each comparison, we extracted the mouse orthologs with e-values smaller than e^−20^. This list of orthologs was then compared to differential expression lists based on RNA-seq analysis of kidneys^47^, tongues^48^, and vaginas^49^ from *C. albicans* infected mice. Gene lists were compared using Venny 2.1.0 (https://bioinfogp.cnb.csic.es/tools/venny/index.html). RNAseq data are available from NCBI’s Short Read Archive (###).

## Results

### *M. sexta* caterpillars are susceptible to the leading fungal pathogen *C. albicans* at 37°C

We first aimed to determine if *M. sexta* fifth instar caterpillars (Fig. 1a), reared and maintained at standard conditions, are indeed susceptible to *C. albicans*. To do so, groups of ten animals were infected with increasing doses of the widely used *C. albicans* laboratory strains SC5314 and SN95^50^. Animals were kept at 25°C, their standard maintenance temperature, and scored for survival on three consecutive days. Dead animals differ from live ones in that their bodies go limp and turn brown-green in colour, which is in stark contrast to the vivid turquoise of live animals (Fig. 1b). Caterpillars infected with *C. albicans* succumbed in a dose-dependent manner. Both *C. albicans* strains killed *M. sexta* efficiently at inocula of 10^6^ or 10^7^ cells per animal (Fig. 1c). To assess if survival measures in caterpillars are comparable to those obtained in the current gold standard, the murine model of systemic candidaemia, we tested *C. albicans* mutants with published phenotypes of either attenuated virulence, such as the *hog1*Δ/Δ^51^ and *ahr1*Δ/Δ^52^ mutants, or wild-type levels of virulence, such as *cka2*Δ/Δ^53^. Cross-species virulence levels are comparable for Hog1, which is just as essential for virulence in caterpillars as it is in mice. Ahr1, while required for virulence in mammals, appears to be dispensable for virulence in caterpillars. Cka2 is not required to establish systemic infections in mammals but is in caterpillars (Fig. 1d).

**Figure 1:**
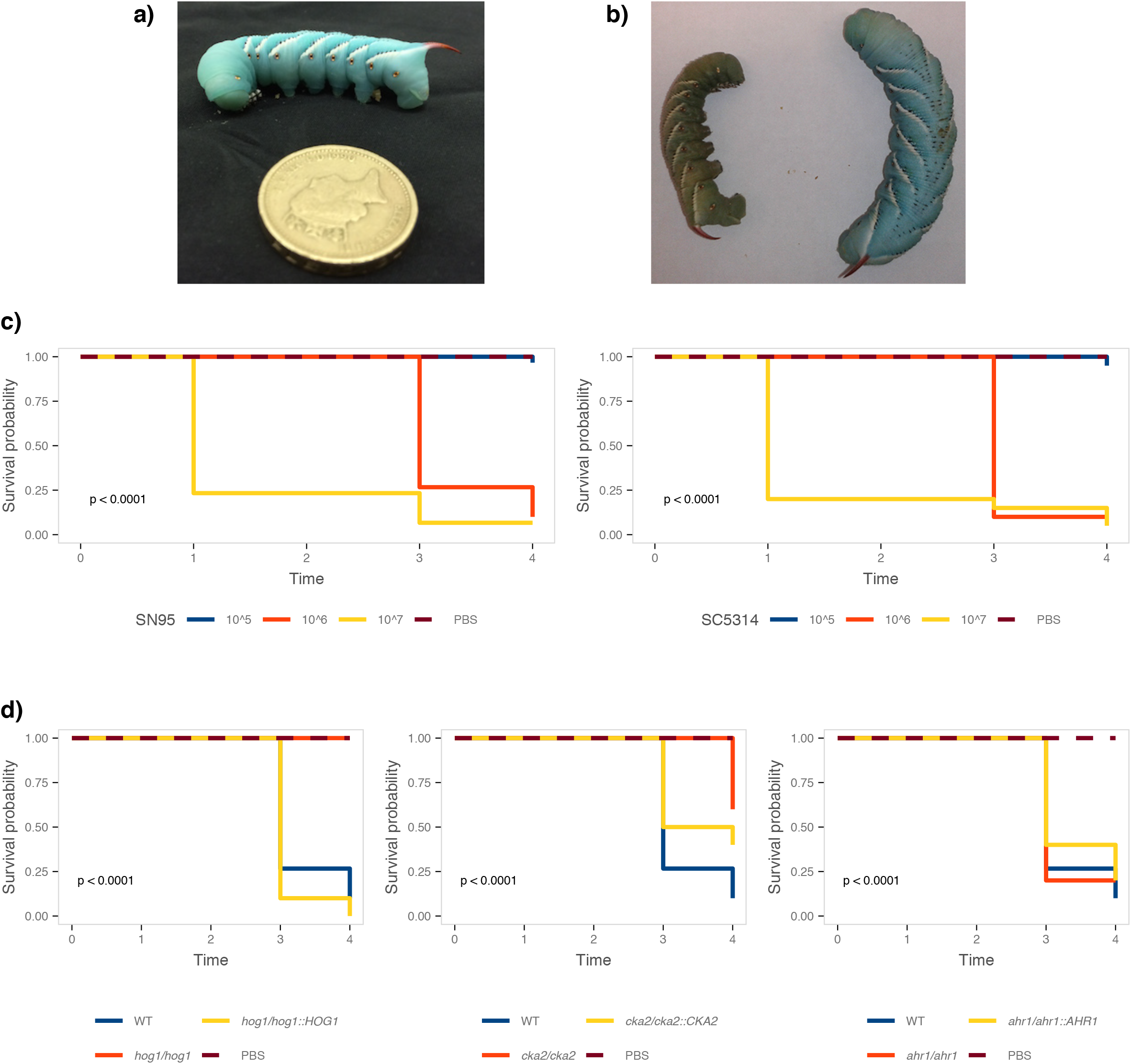
*M. sexta* caterpillars are susceptible to infections with *C. albicans*. **a)** *M. sexta* fifth instar caterpillar prior to injection weighing ~2 g. **b)** 24 hours post injection, the dead animal on the left has lost colour and turgidity compared to the live animal on the right. **c)** Survival curves of animals infected with *C. albicans* SN95 or SC5314. Killing occurs in a dose-dependent manner at 10^6^ and 10^7^ yeast cells per animal. **d)** Survival curves of animals infected with *C. albicans* mutants with attenuated virulence phenotypes in mice or epithelial cell models. The *hog1*Δ/Δ and *cka2*Δ/Δ mutants exhibit attenuated virulence, while virulence of the *ahr1*Δ/Δ mutant is comparable to the wild type. The *hog1*Δ/Δ mutant differs significantly from the wild type (p=0.0001). The complemented strain *hog1/hog1::HOG1* kills *M. sexta* at a level comparable to that of the wild type strain (p=0.21). The *cka2*Δ/Δ mutant is significantly less virulent than the wild type (p=0.00021), while the complemented strain *cka2/cka2::CKA2* is not (p=0.054). Virulence of the *ahr1*Δ/Δ mutant and the complemented strain *ahr1/ahr1::AHR1* does not differ significantly from the wild type (*ahr1*Δ/Δ p=0.72; *ahr1/ahr1::AHR1* p=0.38).

Given the importance of temperature for fungal virulence, we queried if *M. sexta* retained their susceptibility to *C. albicans* at human body temperature of 37°C. Temperature itself does not affect caterpillar survival or development (Fig. S1) but animals are ten times more susceptible to infections with *C. albicans* at 37°C than they are at 25°C (Fig. 2a). At 37°C, 10^6^ *C. albicans* cells per animal lead to 100% mortality on day 4, while 10^7^ cells are required for the same outcome at 25°C (Fig. 1b). To exclude the possibility that mortality is due to starvation rather than the outcome of a host-pathogen interaction, we infected caterpillars with live and heat-killed *C. albicans* wild-type cells and kept them at 37°C. Only live cells, but not heat-killed yeast cells, killed the caterpillars suggesting that killing is not due to nutritional limitations (Fig. S2). Demonstrating susceptibility of *M. sexta* caterpillars to *C. albicans* lent support to them as an alternative host model for the study of fungal virulence and emphasized the need for additional measures of fungal virulence. To add granularity to fungal virulence data collected from *M. sexta*, we complemented measures of survival with quantifications of weight and fungal burden throughout the course of infection. To collect weight data, caterpillars were weighed prior to infection and then daily throughout the course of infection. Weight gain in animals infected with a low dose of 10^4^ cells did not significantly differ from those injected with 1x PBS but caterpillars infected with10^5^ cells per animal exhibited significant weight loss. Too few animals survived infection with 10^6^ cells to allow for a meaningful comparison (Fig. 2b). After establishing susceptibility of *M. sexta* to *C. albicans* at 37°C, we aimed to validate the attenuated virulence phenotype of the *hog1*Δ/Δ mutant strain. To test this, caterpillars were infected with 10^6^ cells per animal with the wild type strain, the *hog1*Δ/Δ deletion mutant, the *hog1/hog1::HOG1* complementation strain and compared to the control group injected with 1x PBS. The *hog1*Δ/Δ mutant is less virulent than the wild type (Fig. 2c). Animals infected with the *hog1*Δ/Δ mutant strain gained significantly more weight than those infected with the wild type strain, while infection with the complemented strain lead to a comparable lack in weight gain (Fig. 2d).

**Figure 2:**
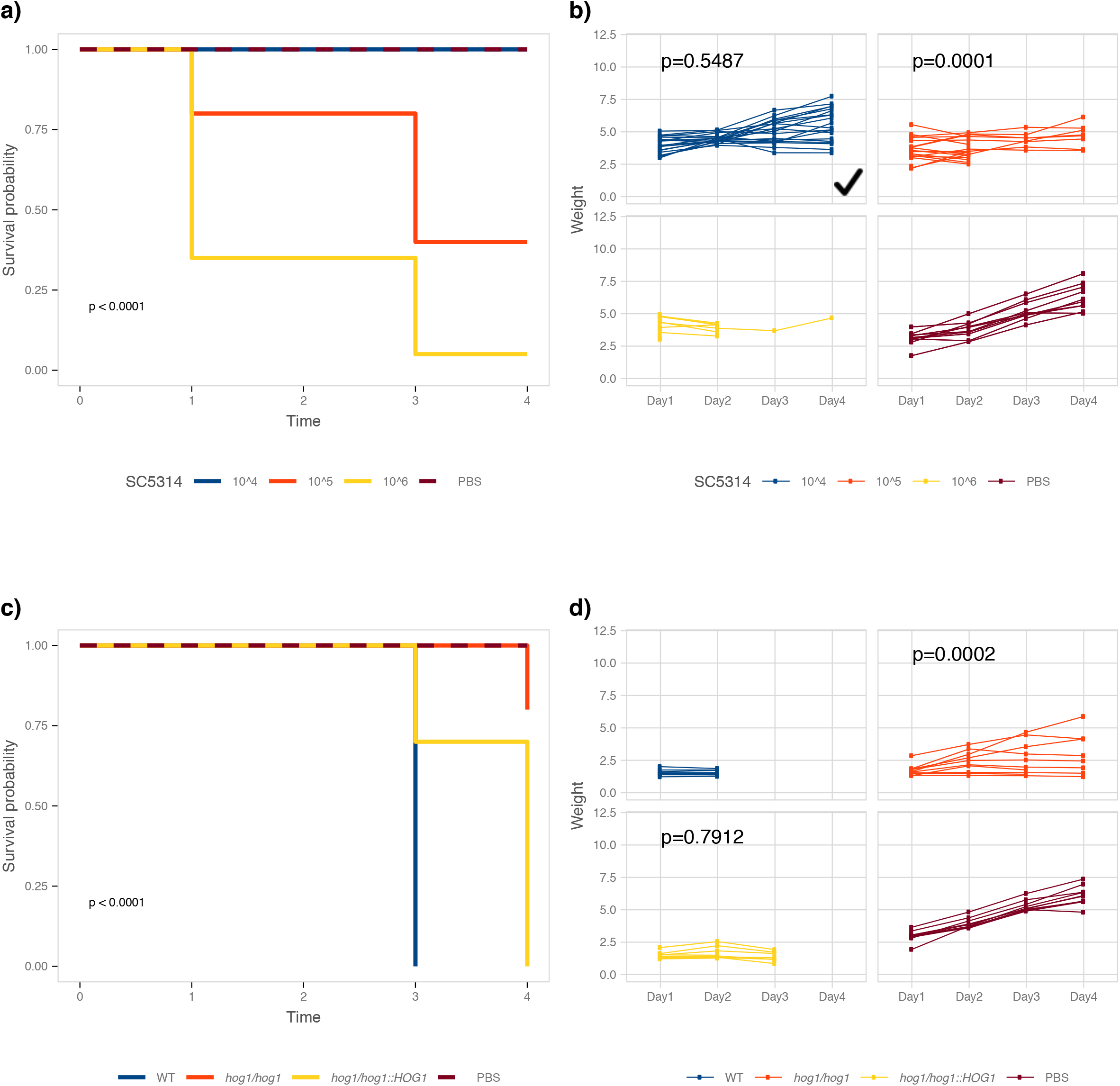
Elevated temperatures increase susceptibility of *M. sexta* to *C. albicans*. **a)** *M. sexta* caterpillars succumb to infection with the laboratory strain SC5314 in a dose-dependent manner at 37°C but less inoculum is required than for infections at 25°C. **b)** The weight of caterpillars infected with 10^4^ cells per animal was comparable to that of caterpillars injected with 1x PBS, while animals infected with 10^5^ cells showed a significant reduction in weight. **c)** Attenuated virulence of the *hog1*Δ/Δ mutant is retained at 37°C. **d)** Animals infected with the JC50 mutant strain gain significantly more weight than those infected with the wild type strain RM1000. Infection with the complemented strain JC52 resulted in a comparable lack of weight gain.

### Measurements of fungal burden in faeces and haemolymph obtained from a single animal throughout infection provides a useful measure of virulence

To further expand the applicability of *M. sexta* caterpillars as a host model for fungal infections, fungal burden in the haemolymph and faeces was quantified daily throughout the course of infection. Since the collection of haemolymph or faeces does not necessitate killing the animal, data could be collected daily throughout the infection for the same caterpillar. Animals infected with the wild type and the *hog1*Δ/Δ mutant were compared to control animals injected with 1x PBS only and fungal burden measured as colony-forming units (CFUs) in faeces and haemolymph in two animals per group. Yeasts were detected in the faeces and haemolymph of animals infected with the wild type on day 2 and CFU counts increased on day 3 (Fig. 3a, b). In animals infected with the *hog1*Δ/Δ mutant, we detected low CFU counts on day 3 in faeces and haemolymph. Even fewer CFUs were detected in the haemolymph of animals injected with PBS on day3. Consistent with the observed increase in fungal burden in wild-type infected animals, indicative of a fungal infection, these animals displayed reduced weight gain while *hog1*Δ/Δ infected animals gained weight at a similar rate as the control animals (Fig. 3c). Thus, measurements of fungal burden, that can be obtained from the same animal throughout the course of infection, provide a valuable parameter for the study of fungal virulence.

**Figure 3:**
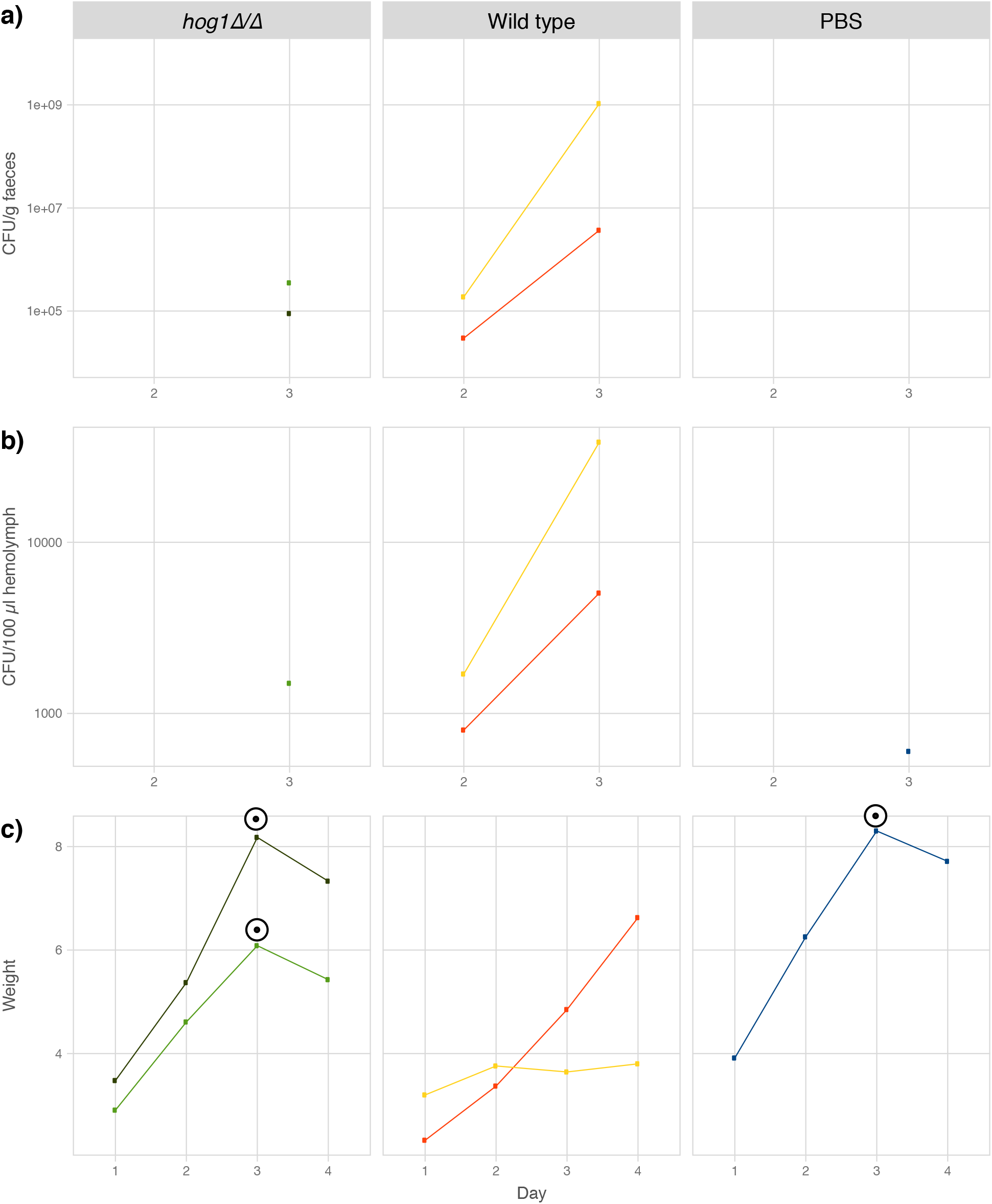
Fungal burden in faeces and haemolymph increases throughout the course of infection. Colony forming units per gram faeces **(a)** and 100 μl haemolymph **(b)** in the *hog1*Δ/Δ mutant (YSD883), the wild type (SN95) and the control animals. Counts increase over time in the wild-type infected animals but not the ones injected with the *hog1*Δ/Δ mutant or PBS. **c)** Weight increases in animals infected with *hog1*Δ/Δ comparable to those injected with PBS, while animals injected with the wild type *C. albicans* strain experience reduced weight gain. The peak and drop in weight observed in the *hog1*Δ/Δ and PBS groups, marked with a ⊙ is coinciding with the onset of pupation. This ‘pupation drop’ is due to the animals refraining from food upon entering the early stages of pupation.

### Caterpillars are suitable for antifungal drug treatment studies

The currently available armamentarium of antifungal drugs is limited and drugs are often lacking in efficacy. The ability to study drug efficacy and mode of action in a host model are thus pertinent to the development of novel antifungal drugs. To determine *M. sexta*’s suitability for drug efficacy testing, we recorded survival and weight of animals infected with the *C. albicans* wild-type strain SC5314 that were treated with increasing doses of two common antifungals, fluconazole and caspofungin (Fig.4). Treatment with fluconazole or caspofungin resulted in overall improved survival and weight gain. Animals treated with 2 mg/kg and 4 mg/kg of caspofungin survived significantly better than those without treatment or those that only received 1 mg/kg of caspofungin. Pair-wise comparisons between different drug concentrations and the untreated control animals yielded no statistical significance for the fluconazole-treated group of caterpillars (Table 2).

**Figure 4:**
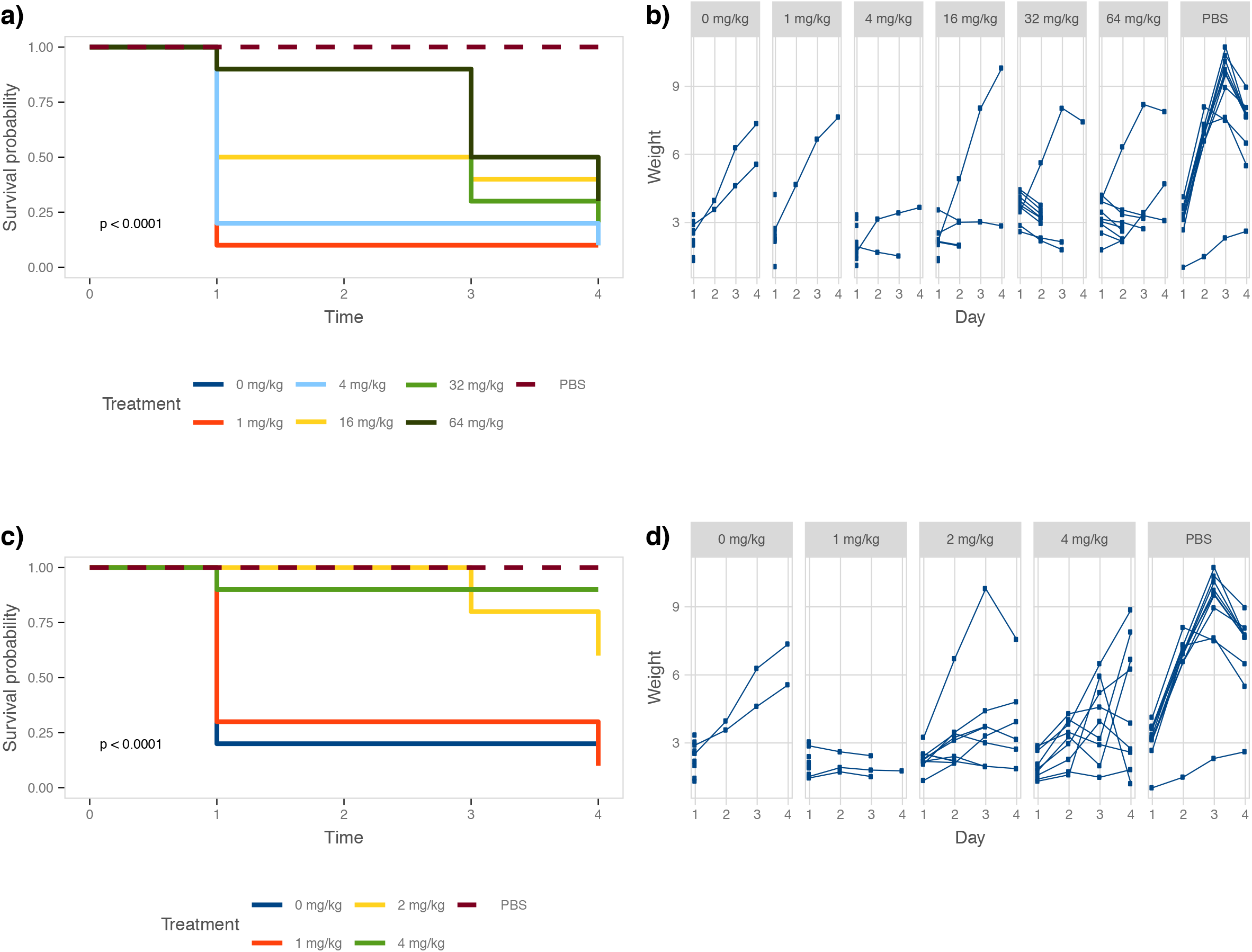
Antifungal efficacy testing of fluconazole and caspofungin. Groups of ten *C. albicans* infected caterpillars were treated with increasing doses of antifungal drug. **a)** Survival improves upon treatment with fluconazole, while weight remains largely stagnant in surviving animals **(b)**. **c)** Caspofungin treatment has a positive effect on *M. sexta* survival and weight **(d)**. Weight data were not analysed for significance due to the lack of surviving animals in the no treatment group.

**Table 2:** Statistical significance values of survival of animals treated with antifungal drugs compared to those not receiving treatment.

| Fluconazole |  | Caspofungin |  |
| --- | --- | --- | --- |
| Concentration | p-value | Concentration | p-value |
| 1 mg/kg | 0.54 | 1 mg/kg | 0.8 |
| 4 mg/kg | 0.63 | 2 mg/kg | 0.015 |
| 16 mg/kg | 0.68 | 4 mg/kg | 0.0022 |
| 32 mg/kg | 0.27 |  |  |
| 64 mg/kg | 0.13 |  |  |

### *M. sexta* caterpillars are broadly susceptible to different yeast pathogens

While *C. albicans* undoubtedly is a leading fungal pathogen of humans, clinical manifestations of fungal infections are not limited to *C. albicans*. To broaden *M. sexta*’s applicability, we sought to quantify the caterpillars’ susceptibility to other fungal pathogens. To this end, animals were also infected with wild type strains of *C. neoformans*, *C. auris*, and *C. glabrata*. Type strains of *Saccharomyces cerevisiae*, the baker’s or brewer’s yeast and *Metschnikowia pulcherrima*, a yeast inhabiting fruits and flowers^54^ served as reference points for attenuated virulence. Animals were infected with increasing doses of yeast cells starting at 10^5^ cells per animal and up to 10^9^ cells per animal. Groups of ten animals per yeast dose and species were then screened for survival and weight daily (Figs. 5 and 6). Notably, only infections with pathogenic yeast species affected survival of caterpillars (Fig. 5). Infections with *S. cerevisiae* or *M. pulcherrima* did not significantly reduce caterpillar survival. Comparing survival amongst the pathogenic yeast species revealed *C. albicans* to be the most severe. 10^7^ *C. albicans* cells per animal result in 100% killing within in 24 hours. 10^9^ *C. auris* cells were required to kill all animals within four days and the same dose of *C. glabrata* resulted in 75% killing. 10^8^ *C. neoformans* cells were required to achieve 50% killing within four days, comparable to *C. glabrata*. It should be noted that due to the high viscosity of the cell suspension, we could not test higher concentrations than the ones stated here. In addition to assessing survival, each surviving caterpillar’s weight was recorded daily. Interestingly, while infections with *S. cerevisiae* or *M. pulcherrima* did not kill caterpillars, infected animals did not gain weight as effectively as uninfected control animals (Fig. 6). Infections with *C. albicans* resulted in severely reduced weight gain, even at the lowest yeast dose tested. Animals infected with *C. auris* and *C. glabrata* responded in a dose-dependent manner, the higher the yeast dose, the more dramatic the weight loss. Notably, animals infected with the lowest dose of *C. neoformans* grew better than PBS-injected control animals. Increasing the yeast dosage, however, resulted in significantly reduced weight gain. This bi-phasic pattern, resembling hormesis^55^ in which a low dose of an environmental agent is beneficial while a high dose is toxic, could be due to *C. neoformans*’ immunogenic capsule^56^. Caterpillars displayed variable degrees of susceptibility to different yeast pathogens and with weight providing an additional measure of host damage. Thus, *M. sexta* caterpillars are broadly applicable for the study of fungal virulence.

**Figure 5:**
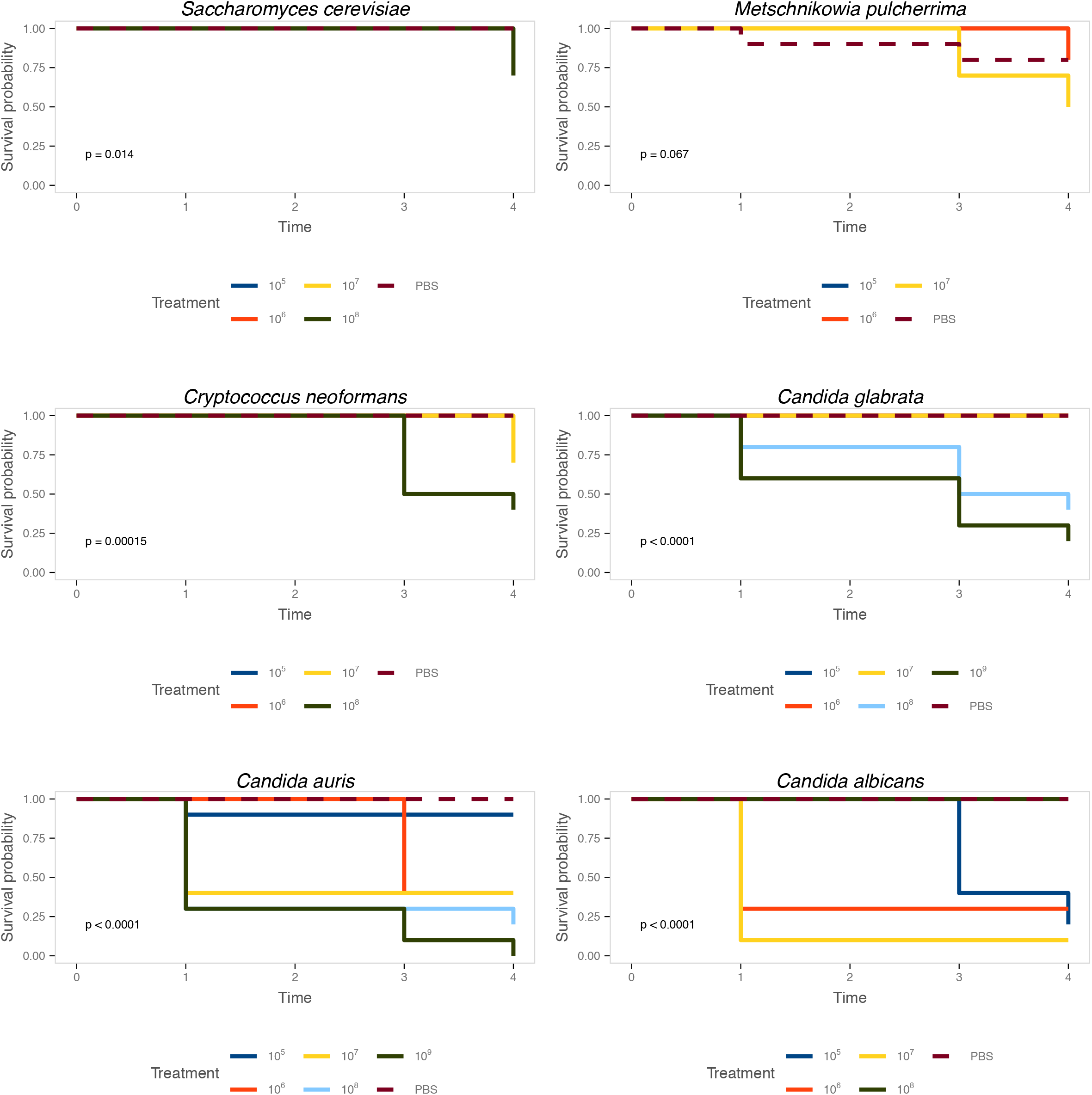
Caterpillars are susceptible to common yeast pathogens. Groups of ten animals were infected with increasing doses of yeasts and survival was recorded daily. Caterpillars are not susceptible to *S. cerevisiae* or *M. pulcherrima* but infections with *C. neoformans*, *C. glabrata*, *C. auris* and *C. albicans* result in significantly reduced survival rates.

**Figure 6:**
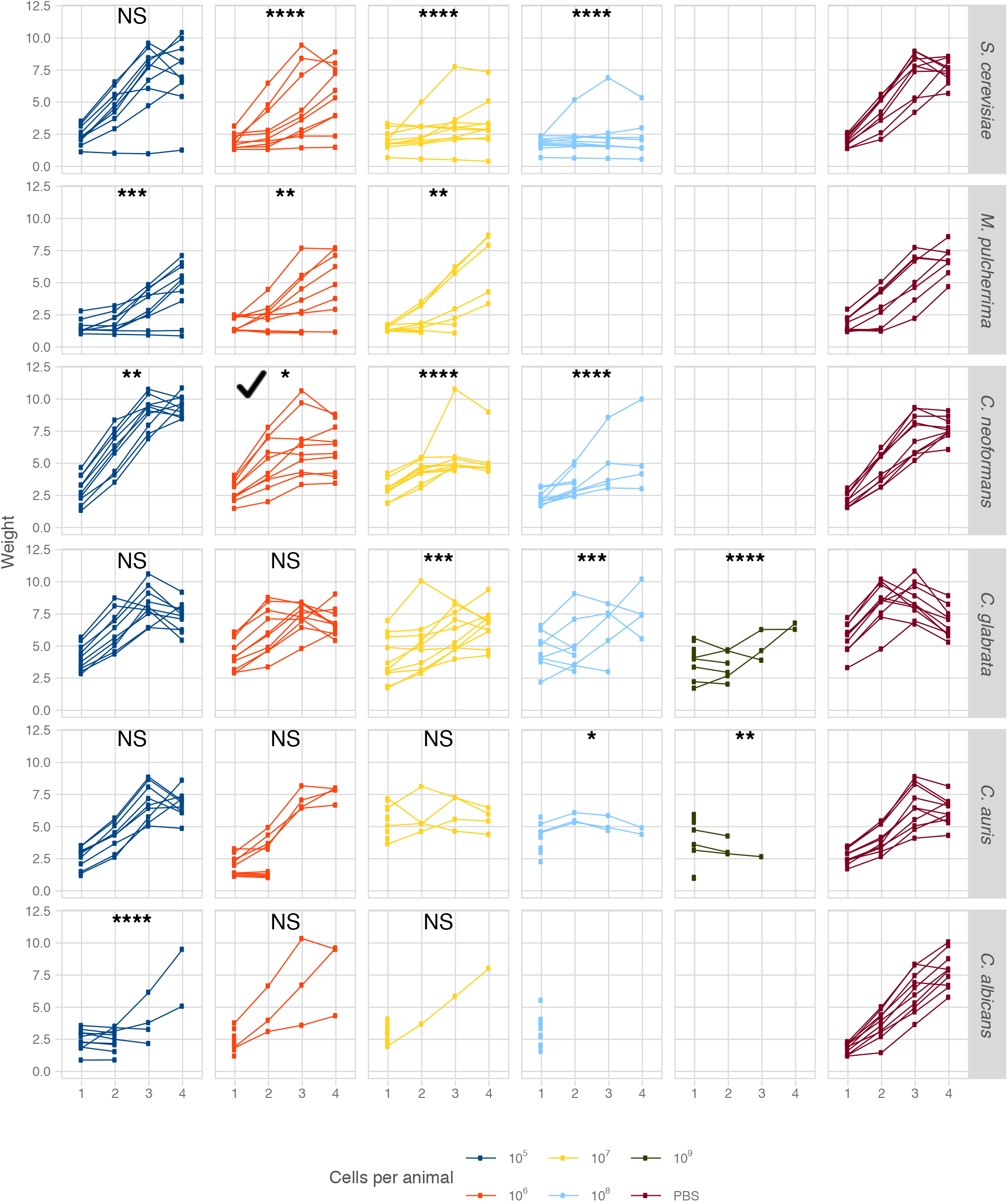
Caterpillar weight as a measure of virulence. Recording caterpillar weight daily revealed that yeast infections affect weight regardless of mortality rates. Daily weight measures were compared to that of PBS-injected animals and statistical significance assessed as follows: * p<0.05, ** p<0.01, *** p<0.001, **** p<0.0001.

### *M. sexta* transcriptional response profiles differ between uninfected and infected animals

As a proof-of-concept we recorded transcriptional profiles in three animals infected with the wild-type strain and compared them to three uninfected control animals. To do so, we collected the mid-gut, rinsed the tissue thoroughly, and extracted RNA, which was then sequenced using Illumina’ HiSeq 2500 platform. With the vast majority of reads mapping to *M. sexta* rather than *C. albicans* (Fig. S3), animals infected with *Candida* displayed a transcriptional profile different from that of uninfected animals (Fig. S4, Table S1). Infection with the wild type strain elicited quantifiable transcriptional changes in the caterpillars. 157 *M. sexta* genes were up-regulated and 165 went down when compared to the PBS control animals (Table S1).

Lastly, we compared our set of differentially expressed *M. sexta* genes to those identified in three murine candidaemia studies. Here, tissue-specific responses in the kidneys^47^, tongues^48^, and vaginas^49^ were recorded for animals infected with *Candida* wild type (Fig. 7). *M. sexta* genes up-regulated during infection and their homologs in all three mouse tissues include serine peptidase inhibitors (Serpinb3c, Serpinb3b), a Ras-like protein (Rit2), transmembrane proteases (Tmprss11a, Tmprss11f), a histidine decarboxylase (Hdc), and a peptidoglycan recognition protein (Pglyrp3). *M. sexta* genes down-regulated in mouse kidney and vagina include a muscle glycogen phosphorylase (Pygm), a bone morphogenetic antagonist of the DAN family (Grem1), meprin 1 alpha (Mep1a), and calponin 1 (Cnn1). Two of these genes are annotated as functioning in immune system processes (Grem1, Pglyrp3) and three more are relevant for signalling and response to different stimuli (Rit2, Pglyrp3, Pygm). Seven genes, however, are involved in protein metabolic processes (Serpinb3c, Serpinb3b, Rit2, Tmprss11a, Tmprss11f, Grem1, Mep1a), suggesting this fundamental process to be of importance in the defence against microbial attacks.

**Figure 7:**
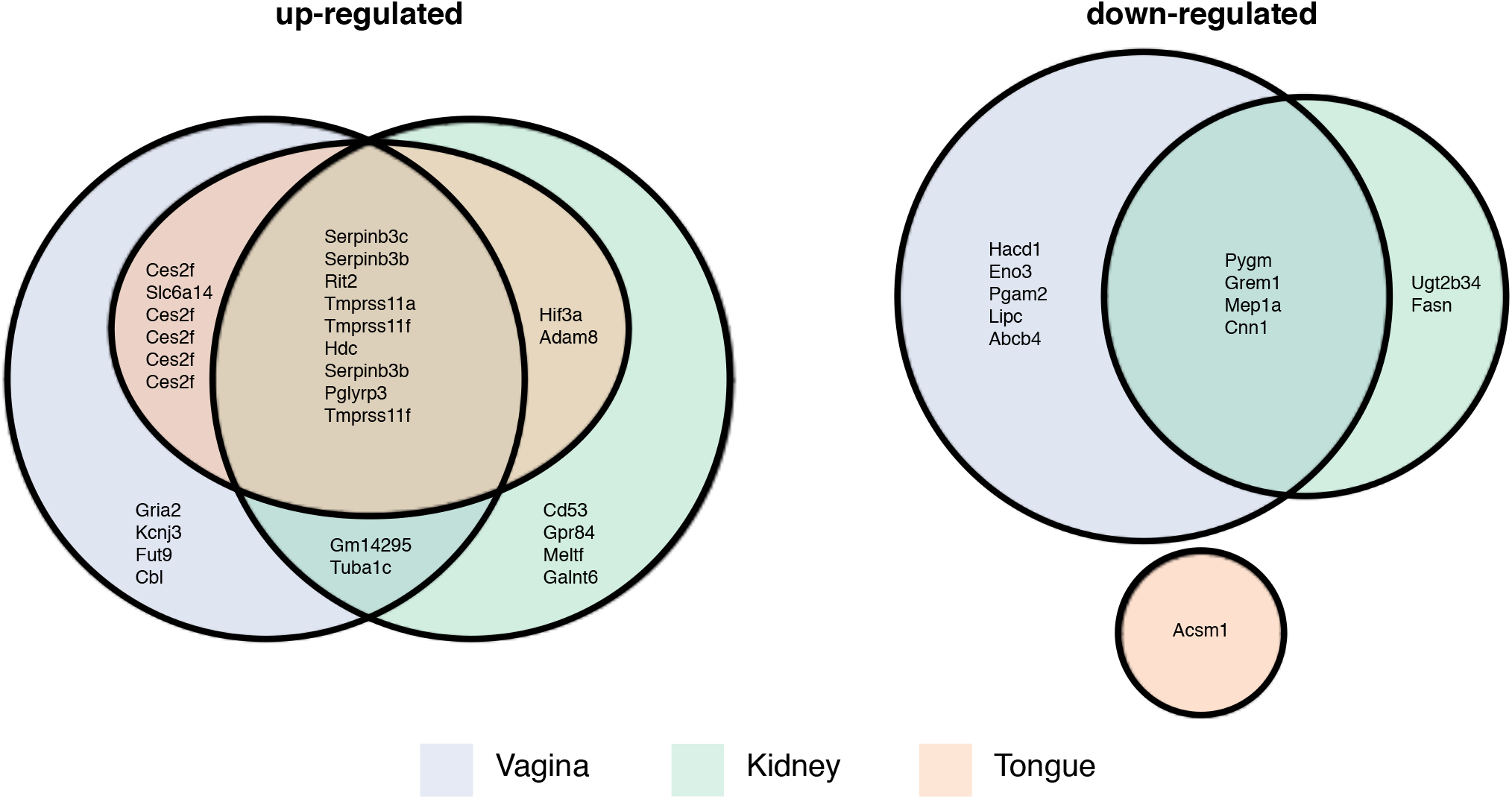
Mouse homologs of *M. sexta* differentially expressed genes. Numbers of *Manduca* genes and their murine homologs that were up- or down-regulated in mouse vagina, kidney or tongue and the overlap between the different data sets as depicted in a Euler diagram. Note, in some cases the same mouse gene was the best BLAST match for several *M. sexta* genes.

## Discussion

Invertebrate host models have become a valuable alternative to mammalian hosts for the study of fungal disease. When choosing the appropriate invertebrate model, care needs to be taken and experimental parameters should be considered^57^. Here we developed *M. sexta* caterpillars as a novel invertebrate model showing that they are naturally susceptible to different human pathogenic yeast species, including the emergent *C. auris*. Unlike other host models, these large caterpillars permit daily measures of fungal burden throughout the course of infection in a single animal by either collecting faeces or haemolymph. *M. sexta* can be maintained at 37°C, is large enough to be injected with a specified yeast inoculum and for weight to be a reliable measure of virulence. *C. albicans* mutant virulence phenotypes found in mice can be replicated and yeast inocula required to elicit a response in caterpillars are comparable to those used in the murine model. Additionally, the caterpillars permit study of antifungal drug efficacy. These parameters commend this invertebrate species as a novel host model for the study of fungal virulence.

This new model system allows for fungal burden to be monitored throughout the course of infection in a single animal via CFU count. This is unlike any other experimental system, where fungal burden is either an endpoint measure in the mouse kidney, in homogenised wax moth larvae^58^, nematodes^59^, flies^60^, or requires genetically modified fluorescent yeast strains for microscopic imaging and analyses^61^. While we did detect very low CFU counts in the PBS control 3 days post infection, we consider this a spurious finding due to cross contamination as preliminary experiments of plating contents of haemolymph and faeces of naïve animals did not detect any yeast growth (data not shown). As a consequence, we amended the protocol to include changing gloves when handling animals of different treatment groups.

When reviewing the weight data collected as part of our study, we noticed two interesting aspects. First, while *S. cerevisiae* and *M. pulcherrima* do not affect caterpillar survival, infections with either species led to reduced weight gain. The dichotomy between survival and weight observed here, further emphasizes that fungal virulence comprises more than a measure of survival. It appears that *M. sexta* would allow discrimination between disease (weight) and death (survival) quantitatively adding further granularity to measuring fungal virulence. Secondly, infections with *C. neoformans* resulted in increased caterpillar weight gain at a low yeast dose but reduced weight gain at higher doses. This pattern resembles the concept of hormesis often deployed by toxicologists to describe the response to toxins, where exposure to a low dose is beneficial while a higher dose results in toxicity due to overcompensation in response to disruption of homeostasis^62^. While tissue-specific hormetic responses have been described in flies, where a virus-acquired cytokine delays ageing^63^, and examples of abiotic factors or signalling molecules affecting peas and aphid infestation^64^, immunity in plants^65,66^, life-span in malaria-transmitting *Anopheles*^67^, and larval development in Black Cutworm^68^, this example here could be the first involving a eukaryotic host-pathogen relationship. The underlying factor remains to be elucidated but *C. neoformans*’ highly immunogenic capsule seems to be an excellent candidate^56^.

Heat-killed *C. albicans* cells appear to be entirely non-pathogenic in *M. sexta* caterpillars as neither survival nor weight are affected by inoculation with heat-killed yeast cells. The lack of mortality in response to inoculation with heat-killed yeast cells indicates that yeast viability and proliferation are required for pathogenesis and excludes the possibility of death due to an allergic reaction in response to a large number of fungal cells. This appears to differ from the responses of other host models to fungal pathogens. While susceptibility was reduced, but still measurable, in *G. mellonella*^69^ and the two-spotted cricket^70^, heat-killed *C. albicans* cells elicited 100% mortality in a sepsis-like murine model^71^. In mice, serum levels of β-(1,3)-glucan levels were elevated in animals injected with heat-killed yeast cells when compared to those infected with live cells. Indeed, heat inactivation leads to increased exposure of β-(1,3)-glucan on the *C. albicans* cell surface^72^ and β-(1,3)-glucan activates the innate immune response in invertebrates and mammals^73^. Given the complexity of receptors involved in recognition of fungal invaders^74^ and the lack of a response in *M. sexta* and other invertebrates indicates that they may not express all or identical receptors as mammals.

Measures of the host transcriptional response provide useful insights into the relevance of specific defence mechanisms. Mouse transcriptional profiles in response to *C. albicans* infections differ between different types of tissues^48,49,75^, which allowed for the identification of tissue-specific response mechanisms such as IL-17 signalling in oral epithelia. Here, we detected numerous differentially expressed genes in the caterpillar midgut when comparing animals infected with *C. albicans* to the PBS control. Given the size of *M. sexta* caterpillars, which permits tissue-specific transcriptional analyses^34^, this phenomenon could be systematically investigated in caterpillars rather than mammalian hosts, who are burdened by ethic concerns and economic challenges prohibiting large scale or time-course studies. Furthermore, partial conservation of the transcriptional response to *C. albicans* infections between insects and mammals offers the opportunity to identify and investigate core eukaryotic response mechanisms.

In summary, *Manduca sexta* caterpillars expand the current repertoire of invertebrate models for the study of fungal disease. They combine a suite of measures that commend it as a new model system. Although, *M. sexta* genomic and transcriptomic analyses are currently still in their infancy, we would expect that the Tobacco Hornworm’s long history of being an invaluable model for diverse facets of biology will lead to reliable tools in combination with genetic tractability and protocols establishing the fungus’ fate inside the caterpillar.

## Acknowledgements

We would like to thank Ewan Basterfield and Chris Apark for expert advice and technical assistance in preparing *M. sexta* caterpillars. Thanks to all the laboratories who kindly shared their strains with us. This work was supported by an ERC Marie Curie Career Integration Grant to SD as well as undergraduate and postgraduate research funding from the University of Bath.

## Disclosure of interest

The authors report no conflict of interest.

## Appendix - Food preparation

Appendix – Food preparation
| Premix |  | Diet cake |  |
| --- | --- | --- | --- |
| Amount | Ingredient | Amount | Ingredient |
| 2,700 g | Wheatgerm | 336 g | Premix |
| 1,260 g | Casein | 1,770 ml | Distilled water |
| 1,080 g | Sucrose | 22.5 g | Agar |
| 540 g | Dried active yeast | 4 ml | Corn oil |
| 360 g | Wesson's Salt | 4 ml | Linseed oil |
| 36 g | Choline chloride | 8 ml | 4% Formaldehyde |
| 72 g | Cholesterol | 0.2 g | Chlorotetracycline |
| 36 g | Methyl paraben | 0.2 g | Vanderssant vitamins |
| 54 g | Sorbic acid | 8 g | Ascorbic acid |

For the premix, deactivate the yeast by microwaving for 5 minutes on low power before mixing all components thoroughly. Store in a cool, dry place.

For the diet cake, heat up 650 ml of water on a hot plate while melting the agar in 1 l of water by microwaving. Combine the agar with the pre-warmed water, 336 g of premix, formaldehyde and oils and mix thoroughly using a stand mixer. Dissolve vitamins, antibiotic, and ascorbic acid in the remaining 30 ml of water and add to mixing bowl once the content has cooled below 50°C to prevent inactivation of vitamins and antibiotic. Line a large ice cube tray with sterile aluminium foil (sterilise by spraying with 70% ethanol) and pour mixture into tray. Let the diet mix set for about 1.5 hours, wrap tightly in aluminium foil and store at 4°C. Keeps for 3 weeks.

## Supplemental Online Material

**Figure S1:**
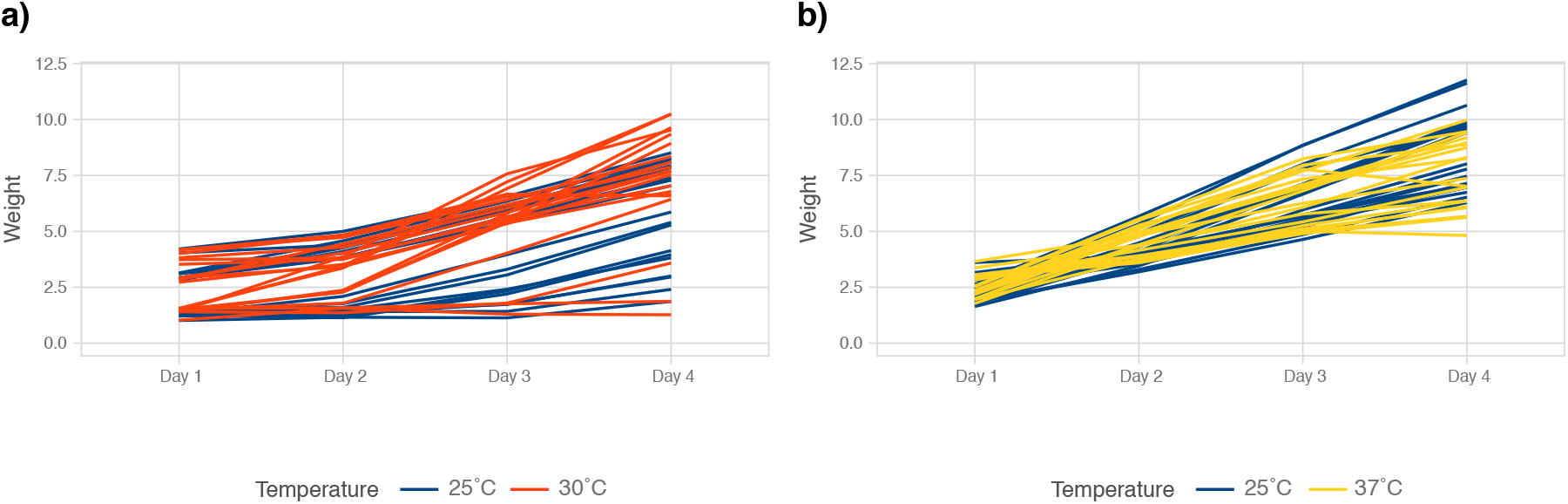
Caterpillar growth is comparable between standard colony temperature and host body temperature. Groups of ten animals were kept on standard diet at 25°C and 30°C **(a)** or 25°C and 37°C **(b)** for four days. Animals were scored daily for survival and weight as a measure of development. Within these parameters, caterpillar development is comparable across all three temperatures.

**Figure S2:**
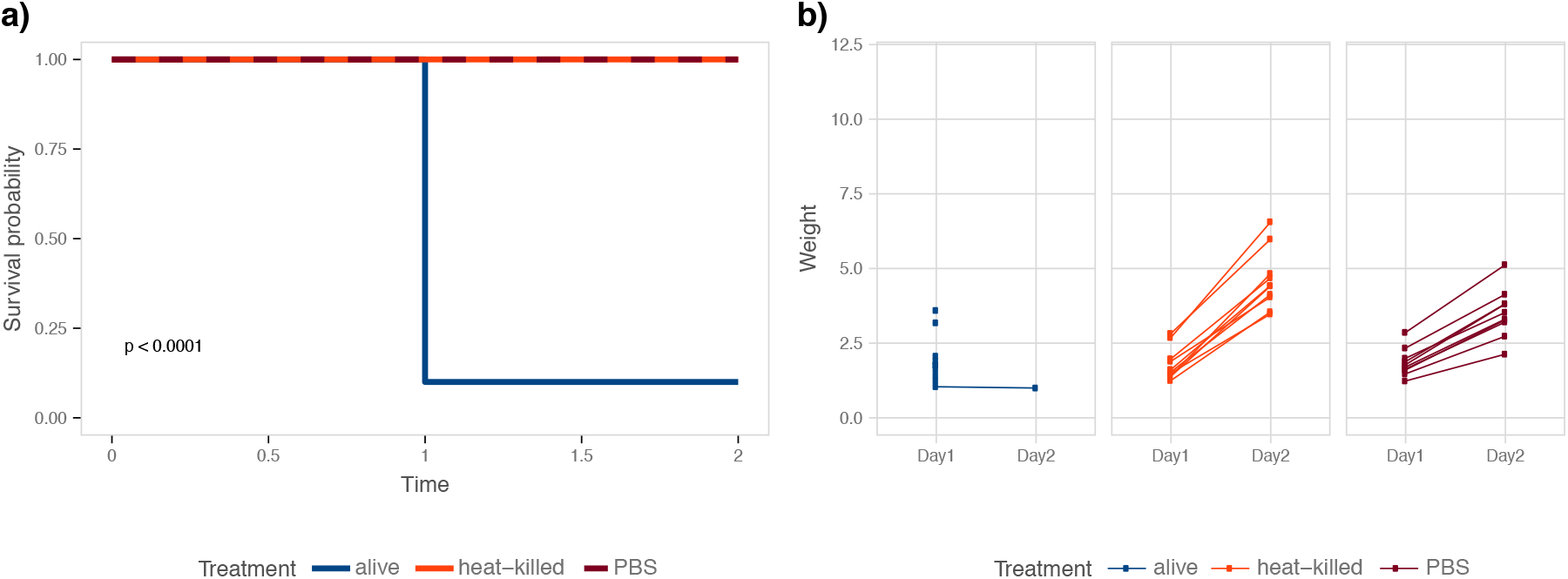
Heat-killed *C. albicans* cells do not affect caterpillar survival or weight gain. Groups of ten animals were injected with either live *C. albicans* wild-type YSD85 cells, heat-killed cells, or PBS. **a)** Only live yeast cells kill caterpillars. **b)** The weight gain in animals infected with heat-killed *Candida* cells is comparable to that of animals injected with PBS.

**Figure S3:**
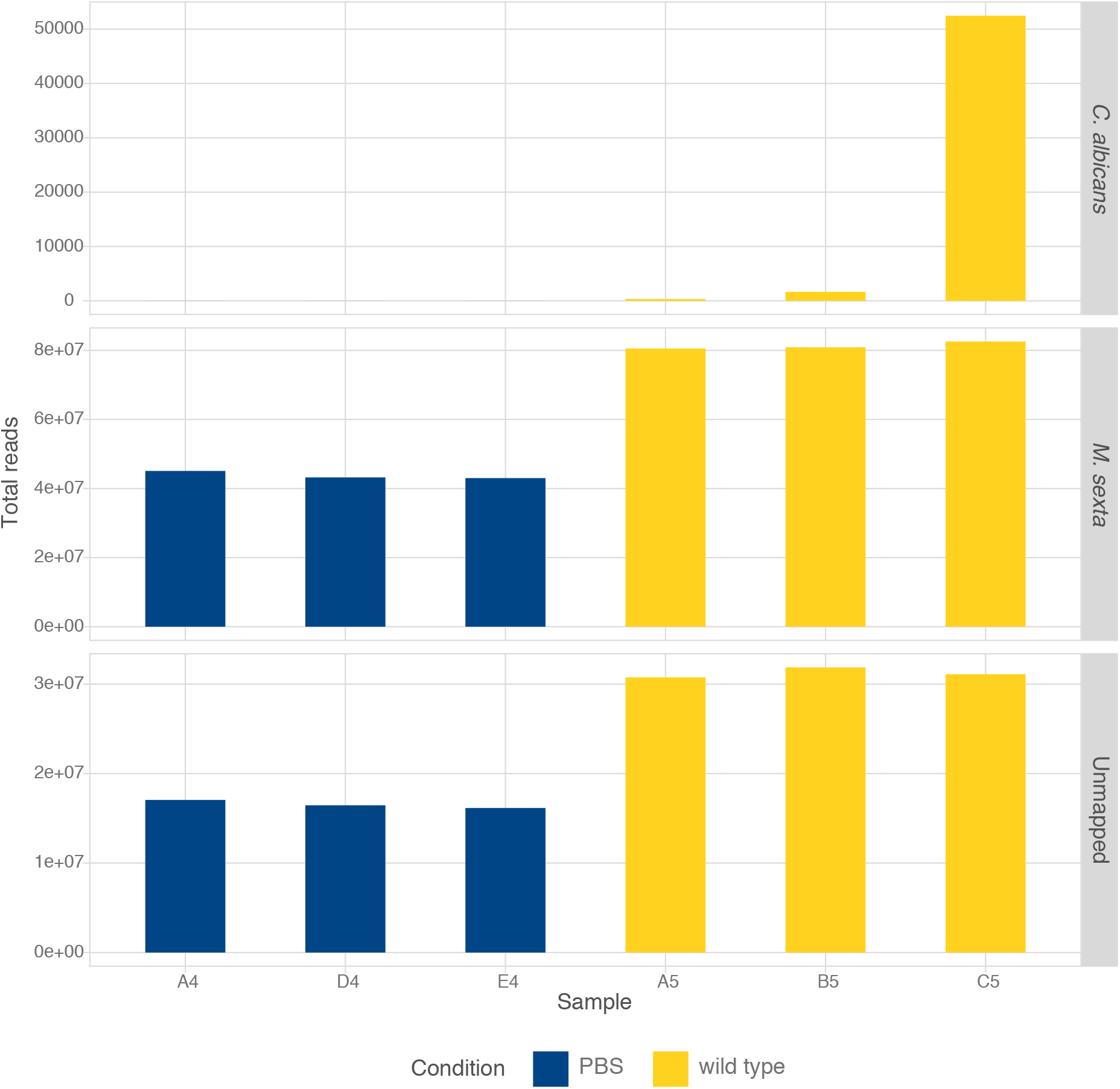
Mapping distribution of RNAseq reads. Between 17.8 M and 34.9 M read pairs per sample mapped either to the *M. sexta* or the *C. albicans*.

**Figure S4:**
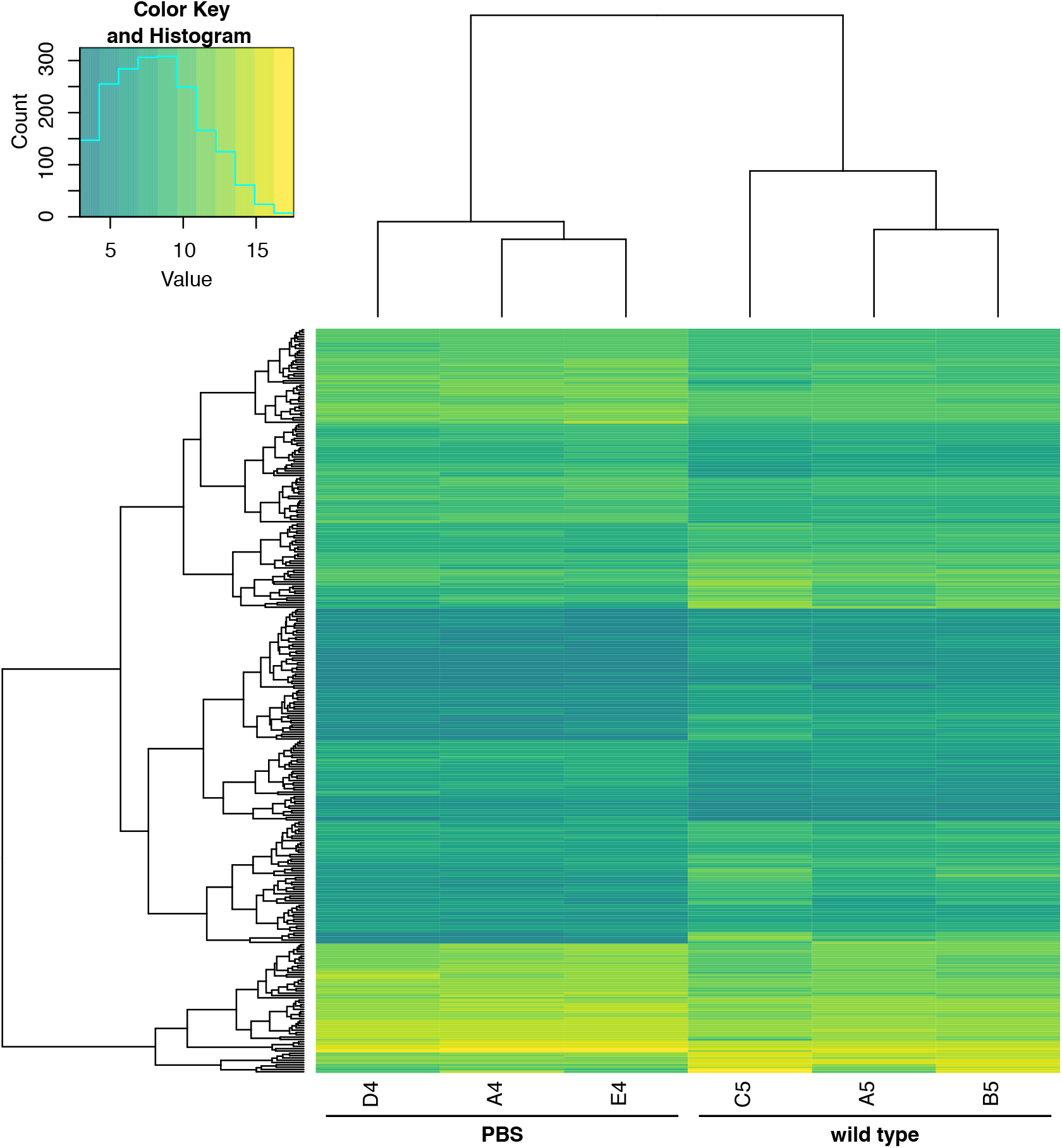
*M. sexta* genomic expression programs in response to infection with *C. albicans*. Hierarchical clustering of global gene expression patterns recorded in samples from uninfected control animals (PBS, samples A4, D4, E4) and those infected with the *C. albicans* wild type (samples A5, B5, C5). Listed are 322 differentially expressed genes on the y-axis in corresponding order to Table S1. In the inset, count refers to the number of differentially expressed genes and value denotes their magnitude of difference.

## Notes

### Competing Interest Statement

The authors have declared no competing interest.

### Summary of Updates

This version contains tissue-specific gene expression data of M. sexta caterpillars infected with Candida albicans and a comparison of these data to expression profiles previously obtained in mouse tissues.

